# Characterization of the pscC (*Type III secretion*) gene of *Pseudomonas aeruginosa* (PA01) and assessment of immunogenicity of pscC protein in rats

**DOI:** 10.1101/071720

**Authors:** SA. Bhuiyan, DJ. Vanitha, H. Sultana, F. Opook, KF. Rodrigues

**Author notes:** **Md Safiul Alam Bhuiyan**, Biotechnology Research institute, University Malaysia Sabah, Malaysia. **Kenneth Francis Rodrigues**, Associate Professor, Biotechnology Research Institute, University Malaysia Sabah, Malaysia. **Daisy Vanitha John**, Associate Professor, Biotechnology Research Institute, University Malaysia Sabah, Malaysia. **Hafiza Sultana**, Assistant professor, Department of Microbiology and Immunology, Rangpur Medical College, Rangpur, Bangladesh. **Fernandes Opook**, Biotechnology Research Institute, University Malaysia Sabah, Malaysia.

## Abstract

Proteins associated with the bacterial membrane can be recruited for application as antigens for the development of vaccines. This preliminary study was directed towards evaluating the antigenic properties of the *Pseudomonas aeruginosa* (PA01) pscC protein which is a component of the Type III secretion system. Gene specific primers were designed to isolate the *pscC* gene which was isolated, ligated onto the multiple cloning site of vector pGS21(a), cloned and expressed in *Escherichia coli* (BL21). The molecular weight of the expressed pscC protein was determined by SDS-PAGE (10% sodium dodecyl sulphate-polyacrylamide gel electrophoresis) and was found to be around 57 KDa and purified by the size exclusion chromatography. Finally, the purified pscC protein was injected subcutaneously into adult Sprague Dawley^®^ rats with a range of concentrations (50, 100 and 150 µg per rat) respectively. Recombinant pscC antigen induced a specific humoral immune response against the antigen, which was validated by Enzyme-linked immunosorbent assay (ELISA). The results concluded that anti-pscC antibody was elicited in the animal model.

## 1. INTRODUCTION

*Pseudomonas aeruginosa* is a versatile Gram negative aerobic bacillus, found in a wide range of environmental habitats. This opportunistic pathogen causes both acute and chronic infections in patients with hospital-acquired pneumonia [1]. It has been classified as the fourth leading cause of nosocomial infection and is associated with cystic fibrosis; burn wound infection, and pneumonic septicaemia [2] [3]. Due to recurrent causes of nosocomial infections *Pseudomonas* infections have become more complex and life-threatening infection, as standard treatments are becoming ultimately ineffective. This organism exhibits intense signs of antibiotic resistance to a wide variety of antimicrobial treatments, including Beta-lactams, Chloramphenicol and Fluoroquinolone [4]. Therefore, specific immune therapy is more desirable than conventional antibiotic therapy [5].

It is an urgent demand of time to implement therapeutic vaccination schemes against *Pseudomonas* infections. *P. aeruginosa* have its own type III secretion system (T3SS), with a protein translocation apparatus and the effector proteins which can be injected into host cells. The secretion and translocation stages comprise over 20 proteins gathered into a needle-like structure termed as injectisome [6]. The T3SS is the part of important virulence determinants of *P. aeruginosa*, in which the pscC protein is a fundamental part of the needle tip for *P. aeruginosa* (Fig 1). The pscC protein has been established to be an important protective antigen of the bacterium and regulate the secretion of translocator proteins to attach with host cell membranes. Once the bacterium interacts with host cell membranes, the T3SS system is activated and this in turn inhibits signal transduction resulting in cellular cytotoxicity or changes in the host immune responses. TheT3SS antigen may react with the host cell, whose maximum component should play a significant role in the host immunity. Therefore, immunization against T3SS antigen can prevent translocon association. However, the functional and regulatory information about this multi-mechanism of pscC antigen will facilitated for the design of novel recombinant vaccines as well as therapeutics.

Effective vaccines are designed to stimulate the innate immune response, as well as carry antigens to specific sub-cellular sites for the elicitation of antigen-specific cytotoxic T cells. Physical delivery to specific locations within a cell is one of the major challenges when developing a suitable candidate for T3SS antigen as a vaccine. Secretory needle antigens of T3SS are easily processed through T3SS pathways. These complexes form a host pathogen interaction is important to identify the molecular pathogenesis for developing an effective vaccine. As a result, the secretory T3SS antigen directly stimulates antigen-specific cytotoxic T cells through the delivery of antigens to the antigen presenting cells (APC), causing a humoral immune response [7] [8]. The proteins for transportation of T3SS pathway have distinct signals that direct them to the secretion machine to stimulate T-helper cells, conferring protection to a diversity of infectious diseases [9].

The preliminary aim of this study was to determine the immune response of the T3SS pscC antigen elicited in animal model. It has been suggested that recombinant or purified protein immunization may allow long-term persistence of immunogenic action in host cells without any risk of infection [10] [11]. Th1 and Th2 cells collectively responses on either Ag are in the secretory product (Th2) or an intracellular or membrane-anchored molecule [12]. These approaches are mostly reliant on the antigen presentation which has been capable to improve immune stimulation to recombinant vaccines. Clinical trials have attested to the safety, efficiency and efficacy, as well as wide application of this immunization technique [13]. The cell surface display of a heterologous antigenic determinant is advantageous for the induction of an antibody against a specific antigen. The orientation of target antigens has been used to develop recombinant vaccines for immunization on rats for the capture and detection of an antibody from serum [14]. The recombinant version of the pscC needle protein of *P. aeruginosa* can be used as a specific antigen for indirect ELISA to screen the antibodies generated against specific protein in response to immunization of the rats.

## 2. MATERIALS AND METHODS

### 2.1 Growth of *Pseudomonas aeruginosa* (PA01) and Isolation of Bacterial DNA

The *Pseudomonas aeruginosa* PA01 culture was obtained from the ATCC. Cultures of P. aeruginosa were grown in autoclaved LB (Luria-Bertani) broth and agar containing peptone (10g/L), Yeast extract (5g/L), NaCl (10g/L) and agar (15g/L). Bacterial DNA isolation was performed via following Aljanabi and Martinez method which was used for rapid salt extraction of high quality genomic DNA for PCR base technique [15]. The plasmid vectors pGS-21a (Gene script, New Jersey, USA) was used as vector for cloning and expression. The complete Freund’s adjuvant (Sigma, USA) was used as an adjuvant for delivery of protein in rats. LB media containing 100μg/ml of ampicillin were used to grow *E.coli* clones at 37°C.

### 2.2 Development of gene constructs for expression of recombinant pscC gene

PCR amplification of pscC gene (1.46kb) of *P. Aeruginosa* was performed using specific primers. Primers were designed according to the pscC gene sequence which was retrieved from the NCBI GenBank (NP_250407). Then nucleotides were selected from both 5’ and 3’ ends of the pscC gene and restriction sites for *EcoR*I and *Hind*III were introduced at the 5’ and 3’ ends respectively in order to facilitate cloning. The forward primer: TCAgaattcCCAGCCTGCCTTACGACTAT, with the restriction site for *EcoR*I, and Reverse primer: CGC-ccatgg-CAACTCGTCGATTTCAAGCA, with the restriction site for *Hind*III. Amplification of the gene was performed 20 µl of total volume containing 2 µl DNA as a template, 4 µl of 5×PCR buffer (Promega), 1.2 µl 1.5Mm Mgcl_2_ (Promega), 0.2 µl of 10 Mm dNTPs (Promega), 0.2 µl of tag polymerase (Fermentas, USA), 2 µl of primer (1st BASE Laboratories, Malaysia) oligonucleotides (Forward & Reverse), and 10.4 µl sterilizes distilled water.

The amplification was performed in a thermal cycle program for initial denaturation at 96°C for 3 min, 30 cycles of 20 Sec at 94°C, 40 Sec at 56°C and 2 min at 72°C. After final extension of 10 min at 72°C, the sample was cool to 4°C. The presence and yield of specific PCR products of approximately 1,500 base-pair was confirmed by gel electrophoresis in 1% agarose gel. DNA samples were loaded in wells and the electrophoresis was carried out at 100 volts. Then the DNA bands were visualized under ultraviolet (UV) trans-illuminator.

### 2.3 Cloning and expression of *pscC* gene

The PCR products were purified using the gel purification kit according to the manufacturer’s instructions (Fermentas, USA). The purified PCR product was cloned into the pGS-21a vector containing T7 promoter to induce the expression of the cloned gene and selected *EcoR*I and *Hind*III were the appropriate restriction sites on pGS-21a vector. Plasmid pGS-21a and pscC gene (PCR products) were further double digested with *EcoR* I and *Hind* III (New England Biolabs, Inc, Beverly, MA). The PCR product of the pscC gene was ligated into the vector using T4 DNA ligase (Fermentas, USA) according to the manufacturer’s instructions. The ligation product was used to transform into *E. coli* of Top10 competent cells by CaCl_2_ method [16]. The insert plasmids were identified by colony PCR. The inserted sequence and its reading frame were confirmed by *EcoR*I and *Hind*III digestion and DNA sequence analysis. The generated pGS-pscC inserted plasmid was again sub-cloned into competent expression cells of *E. coli* BL21 (DE3) following same method described as cloned of Top10 competent cells.

The transformant cells were inoculated into the LB containing ampicillin in test tubes and incubated at 37°C for 16 hours with continuous stirring (200 xg) until the optical cell density reached 0.4-0.6 at OD_600_. Subsequently, the culture was induced with 0.5mM IPTG and maintained at 37°C on a rotary shaker set at 200xg. The un-induced sample (1 ml) was withdrawn prior to induction for used as a control. Afterwards, induced sample was taken hourly and last sample was picked after 16 hours of induction to observe the time course study of the expression. The soluble and insoluble fractions were obtained by treating the pellet with 1ml of PBS (Phosphate Buffer Saline) and lysed using ultra sonication on ice. The insoluble fractions of the protein were run in the 10% SDS-PAGE gel using coomassie Brilliant blue with protein marker (Bio-5150, 1st BASE). After extraction of the intense protein gel, the identification of protein by mass spectrometry was performed. Peptides were extracted according to the Brangan’s methods followed by digestion with trypsin which was analyzed using MALDITOF-TOF mass spectrometer with proteomics Analyzer (48000) [17]. Spectra were analyzed to identify the protein of interest by Mascot sequence matching software with Ludwig NR Database.

### 2.4 Purification and Immunization of pscC protein in rats

The soluble fraction of pscC protein was purified by using by Size exclusion chromatography (SEC). The SEC system (Akta Prime) was calibrated according to the supplier’s instruction. S-100HR column (GE Healthcare) attached to the Akta Prime System was equilibrated using 0.05M Sodium Phosphate, 0.15M NaCl, pH 7.2 at a flow rate of 2.6ml/min. The possible fractions were generated through the significant peaks and collected the specific column for desired protein by chloroform/methanol precipitation method. The fractions were further analysed using 10% SDS-PAGE with Pre-staining Protein Marker.

The male adult Sprague-Dawley Rats (4-8 weeks old) weighing 150-200gms were obtained from Tes Jaya laboratory services, Pulau Pinang, Malaysia. All these animals were acclimatized and quarantined before commencement of the experiment. This study had approval from the Animal Care and Use Committee and Institutional Biosafety Committee (IACUC), Malaysia prior setup the experimental design. Animals were reared in plastic cages using paddy husk bedding at room temperature (25 ± 1°C) and humidity (50 ± 5%) in BSL-1. For evaluation of the immunogenicity of pscC protein, soluble protein was administrated in rats with various doses [Low (50 µg), medium (100µg) and high (150µg)]. Rats were randomly assigned to four groups composed of 6 animals in each. Standard restraints were used during the injection. Group 1 received 100 µl 0.1 M phosphate buffered saline (PBS) with equal of complete Freund’s adjuvant (Sigma, USA) via the subcutaneous route as a control. The Group 2 immunized with 50 µg of total soluble protein mixed with the same volume of complete Freund’s adjuvant. The group 3 & 4 were injected following the same technique with the equivalent of same adjuvant which was treated with 100µg (medium) and 150µg (High) in every 5 weeks. Serum antibody was measured by indirect ELISA and antibody level responds of immunized rat sera which were collected on 7, 14, 21, 28 and 35 days after a single dose of administration. Retro-orbital bleeding was done for blood collection and blood samples were collected under general anaesthesia (95% Diethyl ether with enclosed induction chamber). Serum samples from immunized and control rats were centrifuged at 11,600xg, and stored at −80°C. The collected sera were subjected to enzyme-linked immunosorbent assay (ELISA) assay for the determination of the level of antibodies in sera.

### 2.5 Enzyme-linked immune-sorbent assay (ELISA)

Each sample was screened by indirect ELISA for specific immunoglobulin IgG+IgA (H+L chain specific) using goat anti rat-antibody (Southern Biotech, USA). The recombinant antigen was mixed and poured into the 96-well Polystyrene micro-titre (Membrane solution, USA) plates. After coating the antigenic samples, the micro plates were covered and incubated on a shaker at 4°C for 16hours which was washed three times with 100µl 0.05% (V/V) Tween 20 in PBS, blocked by adding 150 µl of PBS −3%(W/V) BSA followed by incubation for 90min at room temperature. Control and test sera were diluted 1:100 in the blocking agent of PBS-A-0.5% to which 0.05% of Tween 20 was added. Further, after the serial dilution of serum samples at 1:300, 1:600 & 1:900 they were loaded to the micro plate separately and incubated. The plates were washed and applied on goat anti-rat IgM+ IgG antibodies followed by incubation and another wash step. 100 µl of conjugated alkaline phosphatase secondary Abs in PBS–1% and BSA (1/1000) was added to the wells and incubated at room temperature followed by the wash step. P-Nitro phenyl phosphate (Southern Biotech, Birmingham) substrate tablet (5mg/tablet) was dissolved in DSB (1tablet/15ml of DSB) buffer. Subsequently, 100μl of substrate solution was mixed to all tests well and kept at room temperature for 15min until a yellow colour developed. The optical density (OD) was measured using an ELISA plate reader (Infinite ® M-200 PRO, TECAN). The OD results of all samples were calculated as percentage of positive control by using the OD value of the positive control serum by deducting the average OD value of blank serum.

## 3. RESULTS AND DISCUSSION

### 3.1 Plasmid DNA constructions

PCR amplified product was run on a 1% agarose gel and desired band for the respective target pscC gene was found without any primer and dimer. The PCR fragments were found to be approximately 1.46 kb (Figure 2A) amplified from the genomic DNA.

After confirmation, the PCR products were excised out of the agarose gel and purified using a gel extraction kit and digested with restricted enzyme for conformation which was represented in Figure 2B. The synthesized PCR product was then ready for cloning into pGS-21a plasmid vector. The pGS-21a plasmid was constructed by PCR amplification of the coding region of pscC with *EcoR*I and *Hind*III sites and ligating the pGS-21a vector cut with the same enzymes. To determine whether the pscC fragment was successfully ligated in the pGS-21a plasmid, which were then digested with the *EcoR*I and *Hind*III to release the expected size of pscC gene. After 1% agarose gel examination, the RE mixture exposed the presence of two bands. The upper band was consistent to linear according to plasmid size (Approximately 6.2 kb) and another band was corresponding to the expected molecular size of pscC gene around 1.46kb.

### 3.2 Cloning and expression of pscC gene in *E. coli*

The pGS-21a plasmid was transferred to *E. coli* TOP10 and BL21 (D3) and confirmed respectively by restriction digestion, colony PCR and sequencing. The clones were subjected to colony PCR and an amplified 1.46 kb band was obtained as that of positive control. The plasmid showed the presence of 1kb insert and clones were confirmed through PCR and restriction analysis in Figure 3. The PCR positive recombinant clones were also subjected to restriction digestion using *EcoR*I and *Hind*III. The presence of about 1.46kb insert was confirmed by electrophoresis and the positive clone was sequenced with T7 promoter primer (Forward and Reverse). Later than confirmation, clones were further subjected to plasmid extraction, followed by RE and plasmid sequence analysis.

The sequences were then analyzed via BLAST, compared with the reference sequences formerly deposited in NCBI. The sequences was compared to the NCBI database which was revealed about 97% similarity compared the sequences of pscC in BLASTn algorithm. As of the colony PCR the pGS-21a with inserted pscC gene was found approximately 1.46kb size and control showed the 1.0kb. Therefore, it can be concluded that the recombinant plasmid carried the targeted pscC gene.

The protein expression was observed in host strains of *E.coli* BL21 (D3) cells and the most prominent band was appeared at ~84 kDa in 4 samples which was illustrated in Figure 4. The soluble and insoluble fractions were obtained by re-suspending the cell pellet in buffer and sonicated on ice. The slightly visible soluble band for scarce was detected in supernatant phase after visualized under SDS-PAGE (Figure 5). Consequently, a prominent band of the target protein pscC appeared which was visually distinguishable from other proteins based the un-induced controls. Recombinant *E. coli* also showed a distinct band with an approximate cumulative size of ~57 kDa with His tag (6XHis) and GST Tag at the size of 1kDa & 26kDa respectively. For further substantiation, the protein samples were digested by trypsin and peptides were extracted by MALDITOF-TOF mass spectrometer. Spectra were analyzed to identify the protein of interest using Mascot sequence matching software. The result indicated that the amino sequence of pscC protein specified a high ionic scoring match and similar with two peptide sequences appeared in the same protein. The protein identity and mass match were statistically more significant. Protein was analysed and noticed the one matched peptide to the same pscC protein, which was higher confidence for correct protein identified in Figure 6.

### 3.3 Purification and Immunization of pscC protein in rats

The crude fraction was further purified and subjected to gel filtration on a Superdex followed by purification analysis using SDS-PAGE. The chromatogram showed a mixture of monomer and dimer state. The distinct peak was observed in 40 to 47 fractions and the small picks were noticed in 22 to 41 fractions. Additional stage was precipitating the fractions followed by the protocol of Wessel and Flugge [18]. The precipitated proteins were run once again in SDS-PAGE and protein in the lane of fraction 27, 28, 29 columns presented the correct band as their predicted molecular weight of approximately ~84kDa in Figure 7.

After immunization, the antibody levels were measured by indirect ELISA. The result showed that the rat antibody demonstrated a positive value against the immunized pscC protein. As shown in the graphical representation in figure 8, the level of immunoglobulin measured over the 0 to 5th week for all the immunized rats. The OD_405_ Values for different weeks of serum from the post immunized blood were represented in Table 1. The 4th and 5th weeks of all groups of immunized sera were shown in positive value. Four weeks after immunization, The OD_405_ values at 1:300 dilutions were 0.56, 0.64 and 0.63 for the low, medium and high dose respectively. Similarly, at the 5 weeks the antibody level was further increased and the OD_405_ value at 1:300 dilutions to 0.80, 1.04 and 1.22 for the low, medium and high dose respectively. Though for the titer measurement, a standard was needed since we do not have. The OD value of unimmunized and immunized serum was compared in rodent model. It was found that immunized OD value was 2 to 3 folds higher than the control OD value after 3 weeks. The result illustrated that the rat antibody demonstrated a positive value against the immunized serum based on OD value. The levels of the humoral immune responses showed higher antibody of Ag-specific serum immunoglobulin compared to with the controls group. The control groups were always (Group-1) negative to the pscC antigen throughout the study. The humoral immune responses were detected in all plasmid immunized groups and continued increasing after 4th weeks of immunization as indicated by the antibody level. The increased antibody was dose dependant until the 5th weeks. It was noted that the medium dose (100 µg) and higher dose (150 µg) of protein antigen elicited a stronger immune response as compared to the lower dose (50µg).

*Pseudomonas aeruginosa* is a major cause of most opportunistic and nosocomial infections. Although conventional antibiotics are more resistance to *Pseudomonas* organism, now it is challenging to find an alternative immunize therapy. In these circumstances, a monoclonal antibody-based approach is still exploration for the inhibition of *Pseudomonas* infection [19]. Monoclonal antibodies have been established to improve of bacterial permission, stop the colonization and invasion, and avoid the destruction which is triggered by cytotoxic influences [20]. During the past few years, recombinant protein vaccines have shown the highest innovation in modern medical technology. Immunization with microbial antigen has delivered the protective immunity in animal models and considered a potential useful vaccine strategy [21]. Based on previous findings, there are various kinds of immunization have been observed against *Pseudomonas*infection such as killed vaccine [22], purified outer membrane subunit vaccine [23], plasmid immunization [24] and synthetic peptide immunization [25]. The immune response in a mouse animal model to consecutive recombinant protein base vaccine could able to neutralizing the bacterial virulent toxin [26]. It is supported that the use of genetic immunization technique in which a random assortment of genes from a bacteria’s genome to practice for immunization purposes [27]. So, it is supposed to be clear that immunization is the best way of protection against outer membrane secretin antigen of T3SS.

A previous study described about the immunogenicity of *Pseudomonas* cap segment (PcrV) of T3SS needle protein. The anti-PcrV monoclonal antibodies were detected after recombinant PcrV protein was used to immunize in a mouse model. The subgroup of anti-PcrV antibody can repressed T3SS function which was observed in cell lysis assay and designed for assessment of protective activity in a *P. aeruginosa* mouse acute pneumonia [28]. Protection or defence in animal models gives only the recommendation of the potential efficiency of that vaccine in humans [29]. Recently, Mingzi *et al*., 2014 have published the immunogenicity of pcrV gene (T3SS) encoding needle protein was immunized on mice and evaluated the efficiency of vaccines in *Pseudomonas* encoding single antigen [30]. The results revealed that mammalian expression vector with the pcrV gene could elicit a sufficient level of specific antigen to induce the humoral antibody. Based on Brain *et al*., 2001 study, outer membrane of oprF gene of *P. aeruginosa* was immunized by plasmid that indicated that the efficacy of this vaccine can be able to elicited anti-oprF that confers protection of rodents against chronic pulmonary infection. Moreover, the results are similar to previous published results producing defence mechanism in rodent models by immunization with purified outer membrane protein from *P. aeruginosa*[31].

The fact is, without testing its effectiveness in animal models, the recombinant antigen has not yet been applied to induce in the human body for assessment of their efficacy and level of immune responses. It is necessary to conduct a challenging experiment in which live *Pseudomonas* can be injected into immune animals to examine the viability of rat and colony contents of *Pseudomonas* spp. Our further expectation, we tried to determine an applicable vaccine against *Pseudomonas* for clinical trial. Clinical challenge is essential for recombinant antigen efficacy in animal models for assessing their efficacy and level of immune responses. It was also not clear yet, what about the optimal vaccine modalities to induce potent neutralizing antibody responses or to strong humoral immune respond ensures capable to destroy against the infected cell. It is factual that, the recombinant protein vaccine, which is able to induce protection from *Pseudomonas i*nfection, might be more realistic for ending global.

In précis, it is an appropriate challenging strategy for exploration of surface T3SS pscC antigen, which has been characterized from *P. aeruginosa* PAO1 genome as a vaccine candidate. The pscC gene (1.46 kb) encodes a 57-kDa immunogenic outer membrane associated protein of *P. aeruginosa* PA01 strain. The protein was characterized in SDS-PAGE and purified from *E. coli* as a fusion protein to identify anti-pscC specific antibodies in the serum of Sparagury rat. Therefore, it is posited that recombinant pscC protein is capable of becoming a strong carrier and processor for the presentation of target foreign peptides in MHC II to stimulate the T helper cell based humoral immune system in a host cell. The expectations for treatment and prevention of human disease by immunization are changing, and the recurrent refinement of these abilities signify a fit for the current as well as the future manufacture of a vaccine platform against *Pseudomonas* infection.

## 4. CONCLUSION

This study provides preliminary experimental evidence indicating that the outer membrane protein pscC can serve as an elicitor of the humoral immune response in rats. We recommend that future studies focus on a larger sample size and explore a range of open reading frames to determine the precise location of the antigenic component of the pscC protein.

## BIOSAFETY AND ANIMAL ETHICS

The study was approved by the Biotechnology Research Institute Institutional Biosafety Committee (BRIBC). The animal study was carried out in the Level 3 Biological Containment Facility (ABSL-3) at the Biotechnology Research Institute, Universiti Malaysia Sabah.

Recombinant protein expression procedures were carried out using Escherichia coli (BL21) which is classified as a (B) strain that is exempted under the Malaysian Biosafety Law.

